# Dynamics of Adult *Axin2* Cell Lineage Integration in Granule Neurons of the Dentate Gyrus

**DOI:** 10.1101/2023.12.09.570930

**Authors:** Khadijeh A. Sharifi, Faraz Farzad, Sauson Soldozy, Richard J. Price, M. Yashar S. Kalani, Petr Tvrdik

## Abstract

The Wnt pathway plays critical roles in neurogenesis. The expression of *Axin2* is induced by Wnt/β-catenin signaling, making this gene a sensitive indicator of canonical Wnt activity. We employed pulse-chase genetic lineage tracing with the *Axin2-CreERT2* allele to follow the fate of *Axin2*-positive cells in the adult hippocampal formation. We found *Axin2* expressed in astrocytes, neurons and endothelial cells, as well as in the choroid plexus epithelia. Simultaneously with tamoxifen induction of *Axin2* fate mapping, the dividing cells were marked with 5-ethynyl-2’-deoxyuridine (EdU). Tamoxifen induction resulted in significant increase of dentate gyrus granule cells three months later; however, none of these neurons contained EdU signal. Conversely, six months after the tamoxifen/EdU pulse-chase labeling, EdU-positive granule neurons were identified in each animal. Our data imply that *Axin2* is expressed at several different stages of adult granule neuron differentiation and suggest that the process of integration of the adult-born neurons from certain cell lineages may take longer than previously thought.

## 1 Introduction

The adult hippocampus generates new neurons from radial glia-like neural stem cells (NSCs) that are slowly dividing, or quiescent. After activation they proliferate asymmetrically to generate intermediate progenitor cells (IPCs), which differentiate further into neuroblasts. These neuroblasts develop into immature neurons and subsequently into mature granule cells that form synaptic connections and become integrated into the hippocampal circuitry [1].

It is widely accepted that the dentate gyrus undergoes continuous production and integration of granule cells well into adulthood [2]. Postnatally generated granule cells undergo a controlled maturation and differentiation process, followed by integration into existing networks [3]. Although not completely understood, it is thought that immature granule cells work with mature granule cells to integrate sensory stimuli into context, allowing for subsequent contextual discrimination of recurrent similar stimuli [4]. More recent work has allowed for recording of the in vivo activity of immature, adult born granule cells. These younger adult-born granule cells in the hippocampus appear to fire more often but show less spatial tuning specificity than mature granule cells [5]. Conversely, it has been proposed that adult-born granule cells transiently support sparser hippocampal population activity structure for effective mnemonic information processing [6]. The adult-born neurons are also morphologically distinct from neonatally-born neurons [7]. Notably, it is estimated that adult neurogenesis produces half of the granule cells in the mouse dentate gyrus [7].

Several signaling pathways, such as Hedgehog, Notch and Wnt, regulate various facets of neural stem cell (NSC) behavior [8, 9]. Both in the CNS and in other tissues, Wnt signals act as morphogens in a concentration-dependent manner to control progenitor proliferation, tissue domains, and cell fate specification [10]. Moreover, canonical Wnt/β-catenin has a pivotal function in neuronal circuit formation [11].

*Axin2* is one of the first genes induced by canonical Wnt signals, and it has proved to be a valuable genetic tool in the Wnt research field. Wnt/β-catenin-responsive cells have been tracked *in vivo* in many tissues and disease contexts using an *Axin2-CreERT2* allele model [12]. This tamoxifen inducible line has been also used to detect neurogenesis and trace neuronal lineages *in vivo* [13].

However, the duration of controlled maturation and integration processes remains controversial. In 2011, Encinas and coworkers reported that quiescent neural progenitors in the mouse dentate gyrus develop into fully mature cells through multiple controlled and regulated divisions [14]. They showed that this entire maturation process, from quiescent neuronal precursors to mature new cells, takes approximately one month. More recently, Goncalves et al. reported that these new cells form functional connections within 2-3 weeks after their last mitosis, thus integrating into already established networks more quickly than previously reported [15].

In order to further advance our current understanding of the role that the Wnt/β-catenin pathway plays in adult neurogenesis in the dentate gyrus, we conducted a neuronal lineage tracing experiment to measure the proliferation of *Axin2*-positive cells in a pulse-chase manner at various time points after initial tamoxifen pulse. We show that the first evidence of mature granule cells derived from adult neurogenesis is seen 6 months after the initial neuronal stem cell undergoes mitosis. We propose that different cell lineages might have a different pace of integration during adult neurogenesis.

## 2 Material and Methods

### 2.1 Animals and treatments

The *Axin2*^*CreERT2*^ mice [13] were generously received from the Nusse laboratory (Stanford University). The PC::G5-tdT line [16] is maintained in our laboratory. All experiments were carried out in 10-12-week-old mice at the time of induction. All animals were fed standard rodent chow and housed under 12-h light/12-h dark cycle. Four groups of 3 mice were analyzed, for a total of 24 bilateral hippocampal samples. All mice received tamoxifen (100 mg/kg; i.p.) and 5-ethynyl-2’-deoxyuridine (EdU) (40 mg/kg body weight; i.p.) on day t=0. The animals were sacrificed and assessed after 7, 30, 90 and 180 days, based on random assignment to the pulse-chase group. All animal experiments were approved by the IACUC at University of Virginia.

### 2.2 Histology and immunohistochemistry

Mice were transcardially perfused with PBS and 4% phosphate-buffered formaldehyde. Brains were then dissected, post-fixed in 4% PFA and equilibrated in 10% and 30% sucrose, followed by embedding in OCT. Frozen brains were sectioned coronally at 25-μm thickness and stored at - 20°C. For staining, the sections were rehydrated with PBS and permeabilized with 0.1% Triton X-100 (Sigma, USA) for 1 h at room temperature. Next, the slides were incubated for 30 minutes at room temperature and processed with Click-iT™ EdU Alexa Fluor™ 488 Imaging kit (Fisher Scientific), followed by after washes with Triton X-100 in PBS. Next, immunostaining was performed with rabbit anti-Red Fluorescent Protein (RFP, 1:500, Rockland, USA), guinea pig anti-doublecortin (DCX, 1:500, Millipore Sigma, USA); mouse anti-glial fibrillary acidic protein (GFAP, 1:1000, Millipore Sigma, USA), anti-NeuN (1:500, Millipore Sigma, USA), anti-Iba1 (Wako) and anti-Transthyretin (abcam). Secondary antibodies included donkey anti-rabbit Alexa 568 (1:500, Molecular Probes, Invitrogen); goat anti-guinea pig Alexa 647 (1:500, Abcam, ab150187); donkey anti-mouse Alexa 647 (1:500, Molecular Probes, Invitrogen). Slides were mounted with ProLong™ Gold antifade (Invitrogen, USA).

### 2.3 Confocal microscopy and image analysis

Fluorescence signals were imaged with Zeiss LSM-880 with Airyscan confocal microscope (Zeiss, Germany) using sequential scanning mode for Alexa 405, 488, 568 and 647. Stacks of images (1024 x 1024 pixels) were tiled across the dentate gyrus area. Dentate gyri were segmented in the tiled 3D datasets with Imaris 9 (Bitplane, Oxford), and the cells labeled with specific antibodies were detected and counted within the segmented dentate gyrus using the Spots model. Total cell counts in the segmented regions were then normalized per 1000 DAPI-positive nuclei in the dentate gyrus.

### 2.4 Statistical analysis

The results were expressed as Mean ± standard deviation (SD). In all experiments, the statistical significance was set at P < 0.05. Calculations were performed with one-way ANOVA with posthoc Tukey HSD test, or post-test for liner trend, using the statistical package software Prism 6 (GraphPad, San Diego, CA).

## 3 Results

### 3.1 Axin2^CreERT2^ genetically labels neuronal, glial and endothelial cells in the dentate gyrus

To identify *Axin2*-expressing cells in the hippocampus, we crossed *Axin2*^*CreERT2*^ mice [12] to the reporter line expressing tdTomato following Cre recombination (PC::G5-tdT) [16], and analyzed the progeny with immunohistochemistry. Previous reports induced the *Axin2*^*CreERT2*^ allele in embryonic and juvenile mice [13]. Consistent, adult-induced lineage labeled similar cell types in the hippocampus, one week after a single injection of tamoxifen (**Figure 1**). In the granule layer of dentate gyrus, a subset of *Axin2*-positive granule neurons was observed with mature appearing dendrites in the molecular layer. These cells co-labeled with the neuronal marker NeuN (**Figure 1A**). In the sub granular zone, GFAP-positive, ramified cells were observed with a characteristic appearance of astrocytes (**Figure 1B**). *Axin2*-positive cells lacking dendritic morphology were frequently detected in the subgranular zone. Some of these cells co-stained with doublecortin (DCX), a marker of immature neurons [17] (**Figure 1C**). We also show that *Axin2* is expressed in endothelial cells on a subset of blood vessels co-staining with PECAM1/CD31 (**Figure 1D**). Of note, in the third ventricle choroid plexus adjacent to hippocampus, we have also detected *Axin2*-positive cells in the choroid epithelia, but not in myeloid cells (**Figure S1**). Together, *Axin2* is expressed in several cell types in the hippocampal neurovascular unit, showing increased density in the subgranular zone.

**Figure 1.**
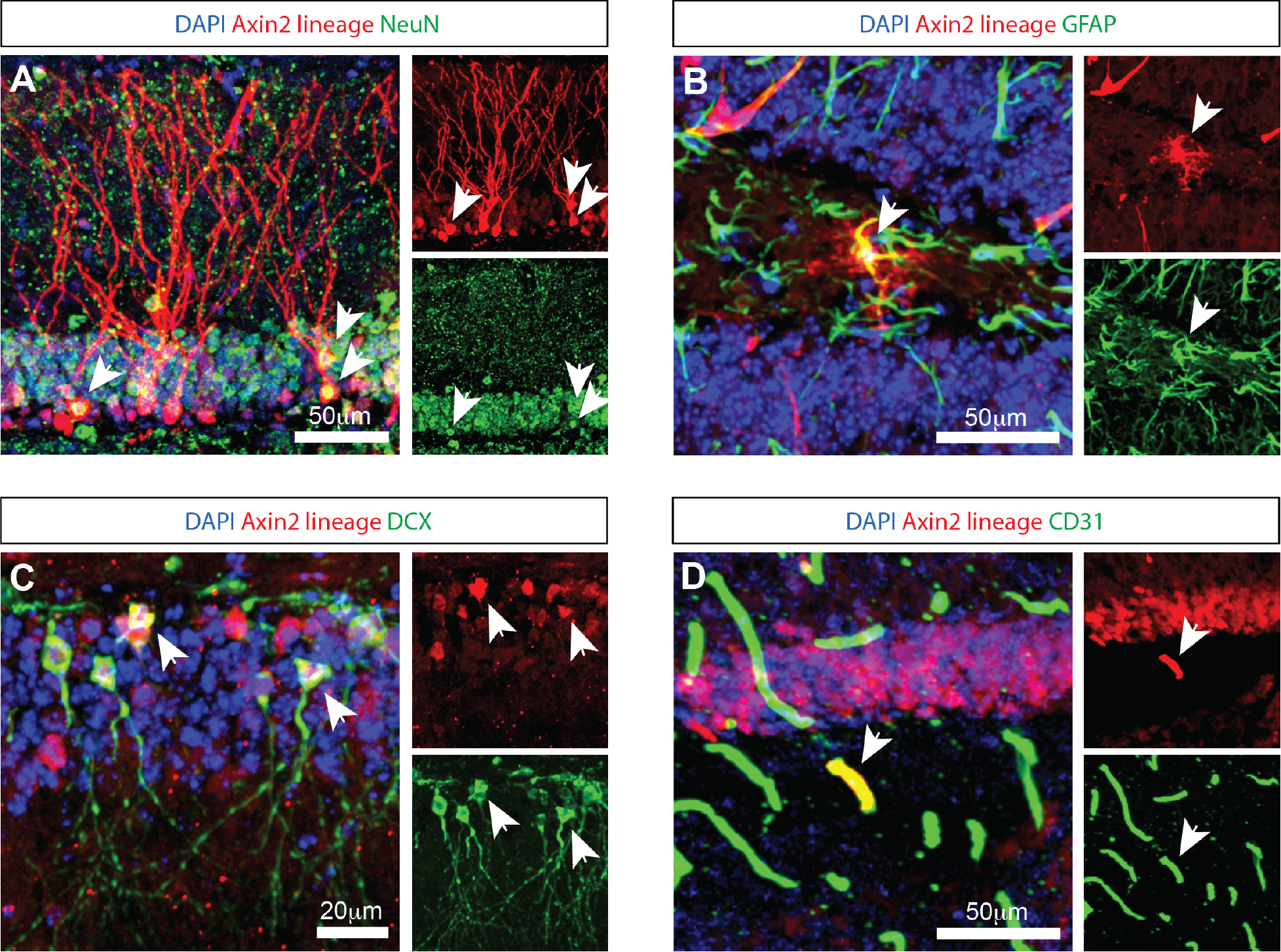
Genetic labeling with *Axin2* ^CreERT2^ reveals neural, astrocytic, and endothelial cells. (A) *Axin2*-labeled tdTomato-positive neurons in the granule layer of the dentate gyrus are NeuN-positive. Arrowheads identify corresponding neuronal cell bodies in the 3-channel overlay (left), as well as in the specific channels showing tdTomato (upper right) and NeuN (lower right). (B) *Axin2*-labeled astrocytes in the dentate gyrus region co-stain with GFAP antibodies. (C) Co-staining with doublecortin (DCX) identifies neuroblasts positive for *Axin2* lineage markers. (D) *Axin2* ^CreERT2^ also labels a subset of endothelial cells in the dentate gyrus region which react with antibodies to CD31. In all panels, white arrows identify double-positive cells. Scale bars, 50 μm, unless indicated otherwise.

### 3.2 Dynamics of the labeled *Axin2* cell population in the granular layer

To follow the fate of *Axin2*-positive cells in dentate gyrus neurogenesis over time, we performed a time course analysis of granule cells labeled with the tracer tdTomato. Adult mice were induced with tamoxifen (100 mg/kg) and followed over time from one week to six months (**Figure 2A**). Differentiated neurons with mature dendrites in the dentate gyrus were counted and normalized per 1000 DAPI-positive nuclei detected with image analysis (**Figure 3 A,E,I;** Methods). Our analysis revealed a statistically significant 6-fold increase (from 2.0 ± 1.1 after 1 week, to 12.0 ± 3.7 after 3 months) in the density of *Axin2* granular cells in the animals sacrificed 90 days after tamoxifen injection when compared to the 1-week time point (**Figure 2K**). There was also a statistically significant positive linear trend for the first three months (p = 0.0432, ANOVA, posttest for trend). The *Axin2*-positive granule cell population peaked around 3 months, but subsequently decreased by 6 months (**Figure 2**). It is also noteworthy that the *Axin2*^*CreERT2*^ line demonstrates baseline leakiness of recombination. As a result, a small number of cells, including granule neurons, become labeled with the tracer even without tamoxifen induction (**Figure S2**). To compensate for this background leakiness, we have measured the number of false-positive granule neurons in three animals and subtracted the normalized and averaged counts from the nominal values determined at all chase time points in this study, as described in the Methods and **Figure S2**.

**Figure 2.**
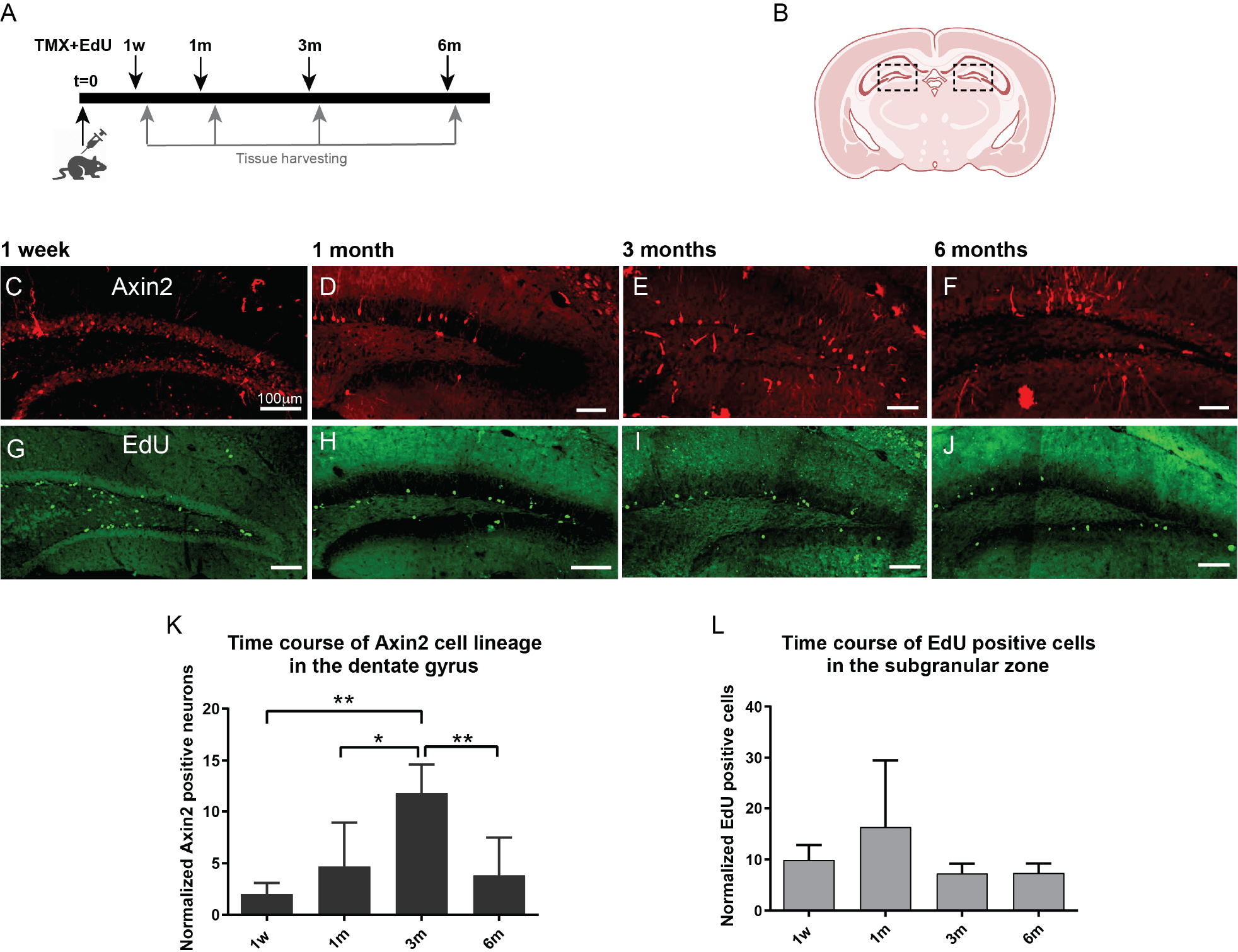
Time course of adult induced *Axin2* cell population in the dentate gyrus. (A) Schematic diagram showing the pulse-chase strategy used in this study. Adult 12-wk-old animals were induced with tamoxifen and concurrently injected with a single dose of EdU. The brains were analyzed at 1 week (1W), 1 month (1M), 3 months (3M) and 6 months (6M) post induction. (B) The central coronal plane used as a landmark for sectioning. The dashed boxes highlight the dentate gyrus (DG) area analyzed in this study. (C-F) Representative confocal images showing tdTomato signal reporting *Axin2* cell lineage. (G-J) Typical EdU staining in DG at different chase time points. The bulk of staining moved from subgranular layer to the deeper granular layer of the DG. (K) Graph showing temporal dynamics of the *Axin2* cell lineage. (L) Time course of EdU-positive cell density during the chase period (1w – 6m). Counts in both plots were normalized per 1000 DAPI positive nuclei. N = 3 mice and 18 sections for each time point. *p < 0.05, **p<0.01. Scale bars, 100 μm.

**Figure 3.**
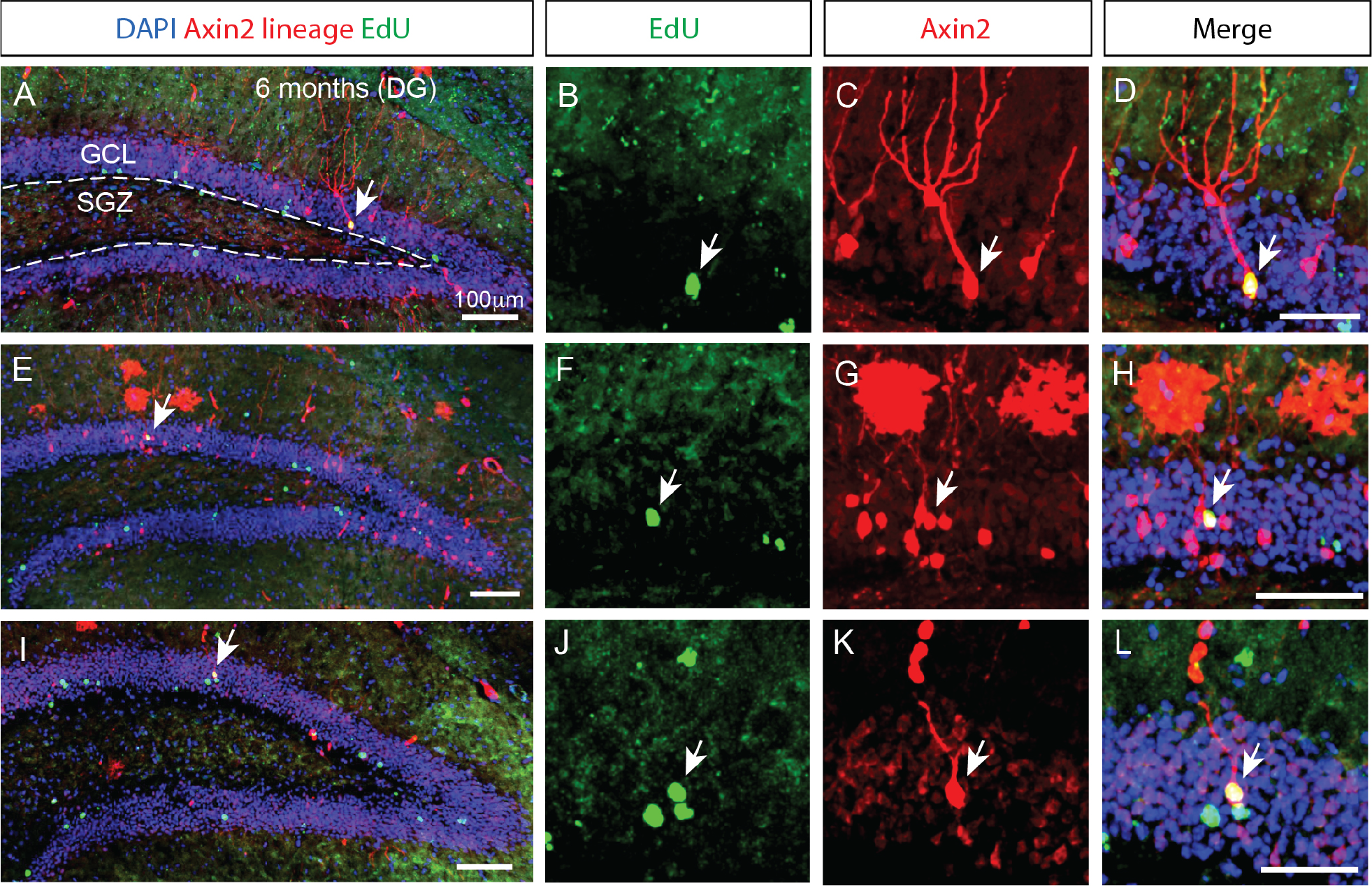
Prolonged integration of mitotically labeled *Axin2*-positive cells in the granular layer of the dentate gyrus. (A, E, I) Low magnification confocal images of dentate gyrus sections from three different animals 6 months after tamoxifen induction. The staining shows *Axin2* cell lineage (red), EdU (green) and DAPI (blue). (A) The granular cell layer (GCL) is outlined with white dashed lines, and the location of the subgranular zone (SGZ) is indicated. Arrows pinpoint the *Axin2*- and EdU-positive cells. (B, F, J) Higher magnification panels showing the EdU signal in the granule cell layer. (C, G, K) Matching panels showing the tdTomato signal indicating *Axin2* cell lineage in the same region as (B, F, J) (D, H, L) Overlay of EdU and *Axin2* panels with DAPI signal, highlighting the mitotically labeled granule neurons originating from the *Axin2* lineage. Low magnification scale bars, 100 μm. High magnification scale bars, 50 μM. DG, dentate gyrus; GCL, granular cell layer; SGZ, subgranular zone.

### 3.3 The time course of aggregate EdU labeling in the dentate gyrus

Next, we investigated cell proliferation in the *Axin2* cell lineage. We performed EdU pulse-chase experiments concurrent with the induction of *Axin2*^*CreERT2*^ labeling. The animals were injected with a single dose of EdU (40 mg/kg, i.p.) and analyzed at the same four time points from one week to six months (**Figure 2 G-J**). Edu-positive cells were detected and measured in the subgranular zone and granular zone, and the values were normalized per DAPI-stained 1000 nuclei (**Figure 2L**). After one month, the number of EdU-labeled cells increased 1.5-fold, but this change was not significant. The numbers of EdU-positive cells remained stable at later time points at approximately 7.3 ± 1.9 EdU-positive nuclei per 1000 DAPI-positive nuclei.

### 3.4 Delayed emergence of EdU-positive *Axin2* granule neurons in the dentate gyrus

While the number of *Axin2*-positive granule neurons peaked three months following the induction, we found no ramified, differentiated neurons containing EdU marker at this stage. However, in the animals analyzed 6 months after the initial pulse, double positive neurons containing the *Axin2* lineage markers as well as the EdU label were detected (**Figure 3**), indicating their derivation from the Wnt-dependent progenitor pool. The neuronal bodies of these double-positive cells were located in the central part of the granular layer (**Figure 3 H and L**). Additional *Axin2*-negative nuclei labeled with EdU were seen in the granular layer (**Figure 3L**), consistent with neuronal identity. Together, our findings confirm that Wnt-dependent *Axin2* cell lineage in the adult brain gives rise to a subset of dentate gyrus granule neurons. Surprisingly, the timeline of maturation and integration of these neurons is considerably longer than what is generally accepted as the time interval needed for neuron differentiation during the adult neurogenesis (**Figure 4**).

**Figure 4.**
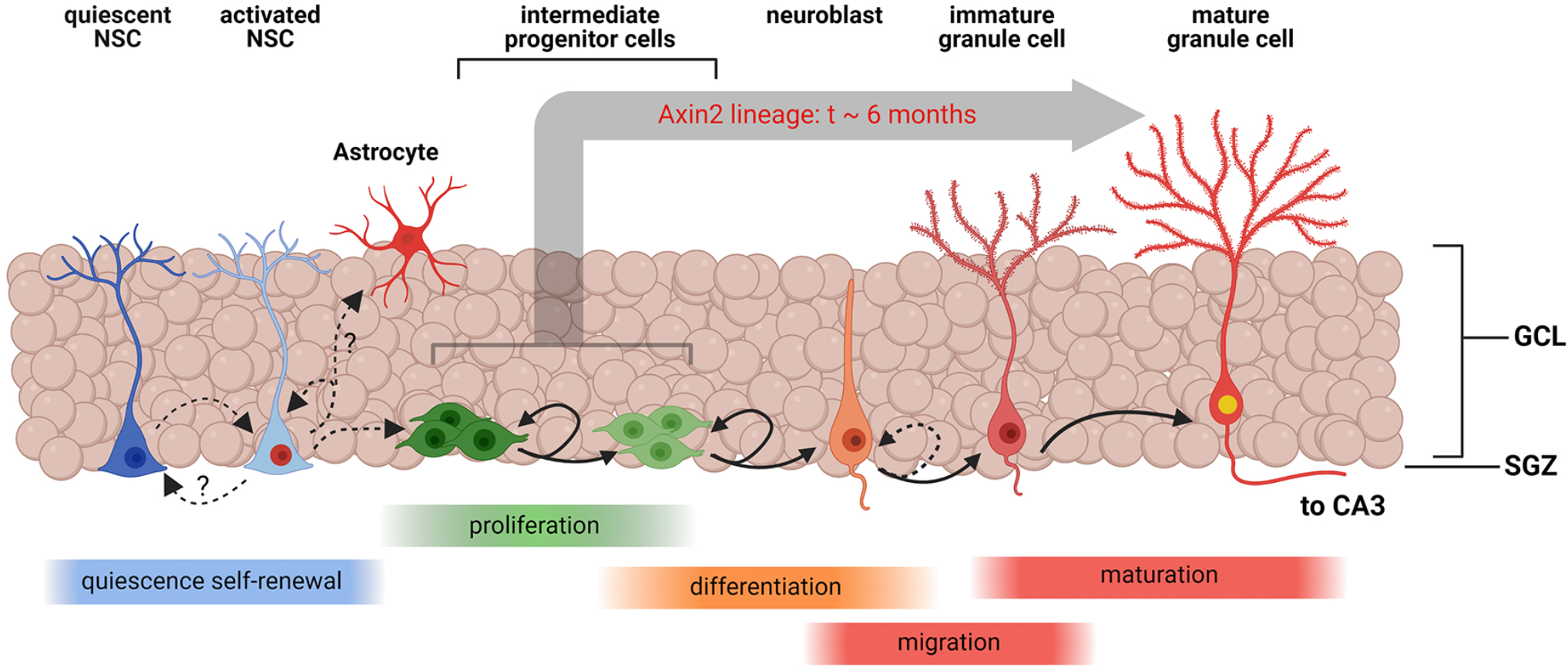
Schematic representation of Wnt-associated subgranular neurogenesis. A summary diagram of the stage in subgranular neurogenesis from quiescent neuronal stem cells (NSCs) to mature granule cells. Quiescent NSCs are multipotent undifferentiated cells. Once induced, quiescent NSCs differentiate into activated NSCs, which may replenish the pool of quiescent NSCs, or commit to the astrocyte maturation pathway. Activated NSCs also develop into multipolar intermediate progenitor cells and neuroblasts which continue to proliferate and eventually migrate inside the granular layer and enter the granule cell maturation pathway. *Axin2*-positive NSCs and progenitors represent a sub-lineage of the cell lines, contributing to subgranular neurogenesis. In this study, we have unequivocally detected *Axin2* lineage markers in the intermediate progenitors and neuroblasts, which are amenable to mitotic labeling, and post-mitotic mature neurons. Our data suggest that the process of integration of progenitors into mature granule cells can take as long as six months in this cell lineage.

## 4 Discussion

We have characterized the cell fate of the adult induced *Axin2* cell lineage. While others have previously shown that *Axin2* expression is active in hippocampus [13], the genetic labeling in their study was induced at the embryonic and juvenile stages within first two postnatal weeks of development. Our experiments confirm that *Axin2* plays role in the neurogenesis of granule cells of the adult dentate gyrus. Consistent, *Axin2* labeling was observed in DCX-positive immature neurons. Conversely, we have not identified *Axin2* in NSC or IPC cells, and more precise characterization of the *Axin2* progenitor population is warranted. *Axin2* expression has been previously reported in astrocytes [18]. However, it is also possible that tdTomato-labeled astrocytes are born from the *Axin2*-positive NSCs (**Figure 4**), and thus derived from the *Axin2* domain in NSC. We have noticed larger counts of tdTomato-labeled astrocytes at 3- and 6-month pulse-chase time points, but these increases were not formally analyzed. Furthermore, we have also shown that *Axin2* expression is present in microvascular endothelia, in line with the role of Wnt/β-catenin signaling in angiogenesis and vascular integrity [19-22]. Relatedly, *Axin2* is also expressed in the choroid plexus. We found that the *Axin2* cell lineage did not contain any myeloid cells, neither microglia nor macrophages. Instead, these cells co-stained with Transthyretin, a marker of choroid plexus epithelial cells (**Figure S1 E-H**), [23]. The choroid plexus generates cerebrospinal fluid, and the epithelia distributes thyroid hormone and secretes numerous factors, including Wnt proteins. Thus, the *Axin2*-lineage of choroid plexus can potentially affect cerebral proliferative processes in a long-range manner [23, 24].

The *Axin2* granule neurons presented a dynamic cell population in our experiments. Our measurements imply that the *Axin2* positive granule cell population is replenished from the *Axin2*-labeled progenitors and increase six-fold after three months. Interestingly, our work also showed that the *Axin2* positive granule cells subsequently decreased between 3 months and 6 months. We believe that this could indicate a more limited lifespan of these neurons. However, further research is necessary to address this possibility.

Encinas et al have demonstrated that both NSCs and amplifying IPCs decrease with age in mouse lines [14]. Yet here we show that the *Axin2* arm of the canonical Wnt pathway remains active in the adult dentate gyrus neurogenic niche. Along with other neurogenic factors, like sonic hedgehog (Shh), bone morphogenetic proteins (BMPs), and Notch, Wnt ligands are known regulators of adult hippocampal neurogenesis [11, 25]. Wnts are expressed by local astrocytes and NSCs themselves, acting as paracrine and autocrine factors in the NSC niche [26, 27]. Specific ligands such as Wnt3 have been shown to promote neuroblast proliferation and neuronal differentiation through the canonical Wnt pathway [26]. In addition, Wnt signaling also emerged as important pathway promoting multipotency and self-renewal of NSCs [28]. However, the specific roles of *Axin2* in these processes remain to be characterized.

One important question to address is whether different subsets of NSCs exist with various degrees of quiescence and distinct abilities to generate neurons and glial cells. Genetic lineage tracing with inducible version of the Cre recombinase under different promoters has provided evidence for the functional heterogeneity of NSCs in the dentate gyrus [29, 30]. Some NSCs have been shown to be neurogenic and short lived, while others are multipotent and self-renew for longer periods [31]. Using genetic labeling with nestin-CreER and performing a double pulse-chase experiment with tamoxifen and BrdU, Encinas detected BrdU- and NeuN-positive granule neurons in the nestin cell lineage 30 days after pulse-chase induction [14]. This is in stark contrast with our results suggesting that *Axin2* cell lineage requires much longer for neuronal differentiation and integration into granule neurons. Since *Axin2*^*CreERT2*^ labels fewer neuronal progenitors than nestin-CreER, some co-labeled cells could have been missed. It is surprising, however, that in comparison to the 6-month time point, twice as many *Axin2*-positive cells were seen after the 3-month pulse-chase, with a similar general level of EdU staining, yet no tdTomato+ EdU+ neurons were detected (Figure 3 K, L).

We acknowledge that the labeling approach used in this study was rather sparse, consisting of a single simultaneous injection of tamoxifen and EdU. Designed to facilitate effective lineage tracing, the resulting labeling density did not provide sufficient labeling of stem cells and neuronal progenitors, as stated above. More saturating labeling strategies will have to be employed in future studies to reveal the presence of *Axin2* in quiescent adult neural stem cells

## 5 Conclusions

In summary, our data supports the view that Wnt-responding and *Axin2*-positive progenitor cell population contributes to adult dentate gyrus neurogenesis. We conclude that integration of *Axin2*-positive progenitors may take considerably longer than reported for other cell lineages. This outcome warrants further investigation of the *Axin2* sub-lineage compared to other granule neurons.

## Acknowledgements

This work was supported with Focused Ultrasound Foundation (FUSF) seed grants and NIH R21NS116431 funding to P. Tvrdik. We thank Drs. Kevin Lee and Elise Cope, Department of Neuroscience, University of Virginia, for critical reading of the manuscript and valuable comments.

## Conflicts of Interest

The authors declare that there is no conflict of interests regarding the publication of this paper.

## Author Contributions

M.Y.S.K, K.A.S. and P.T. designed research, K.A.S. carried out the surgical, fluorescence and biochemical experiments, K.A.S. and F.F. analyzed the data. M.Y.S.K provided resources and participated in conceptualization, K.A.S. and P.T. designed genetic crosses for the experiments. K.A.S, F.F. and P.T. wrote the paper, P.T. directed the project, edited the manuscript.

## Data Availability

Data supporting the findings in this study are available from the corresponding authors upon reasonable request.

## Notes

### Competing Interest Statement

The authors have declared no competing interest.

