## Supplemental Material for "Dynamics of Adult *Axin2* Cell Lineage Integration in Granule Neurons of the Dentate Gyrus"

**Running Title:** Wnt-dependent adult neurogenesis

*To whom correspondence should be addressed:

Petr Tvrdik, PhD, Department of Neurosurgery and Neuroscience, University of Virginia Health System, Charlottesville, VA.

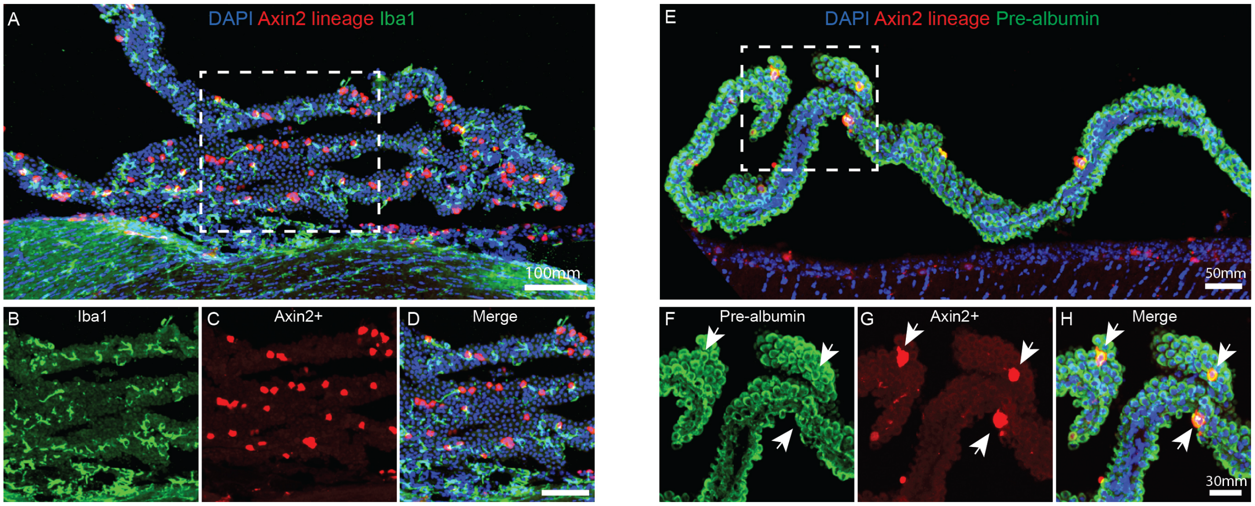


**Figure S1 Confocal analysis of *Axin2* expression in the choroid plexus.**
**(**A) Section through the choroid plexus from the 3rd ventricle one week after tamoxifen induction. The section was stained with anti-Iba1 antibodies and nuclear stain DAPI. Scale bar, 100 µm.
(B) Zoomed region, framed with the white dashed box in A, showing the Iba1 signal.
(C) The same field of view showing direct fluorescence of the *Axin2* lineage marker tdTomato.
(D) Overlay demonstrating that Iba1 and tdTomato positive cells are mutually exclusive. Scale bar, 50 µm.
(E) A coronal choroid plexus section from same animal incubated with anti-Transthyretin (Prealbumin) and DAPI. Scale bar, 50 µm.
(F) Zoomed-in detail showing anti-Transthyretin staining.
(G) A corresponding field of view showing direct fluorescence of *Axin2* - tdTomato.
(H) Overlay demonstrating that Axin2-tdTomato cells co-label with Transthyretin (Prealbumin), (denoted with white arrows). Scale bar, 30 µm.

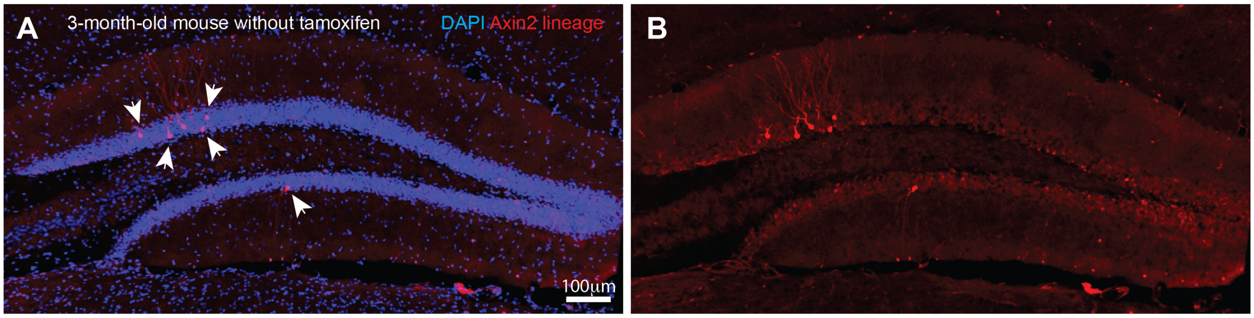


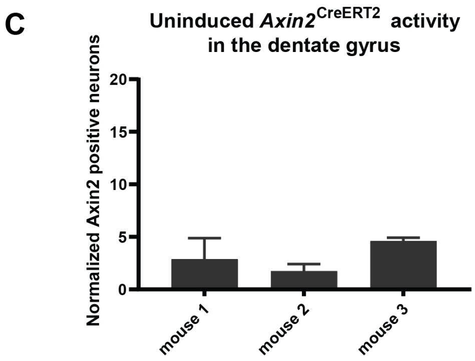


**Figure S2. Uninduced *Axin2* $^{\mathrm{CreERT2}}$ activity in the adult hippocampus.**
(A) Representative confocal image of the dentate gyrus (DG) showing *Axin2* cell lineage (red) and DAPI (blue) staining in the brain sections from 3-month-old animals that were not induced with tamoxifen. The arrows point out the *Axin2*-positive cells.
(B) A corresponding single-channel image showing the tdTomato signal only, demonstrating *Axin2* cell morphology consistent with neuronal differentiation.
(C) Graph showing the average density of *Axin2* positive cells from 3-7 DG sections in 3 different animals (N = 13) without tamoxifen induction. These counts, normalized per 1000 DAPI positive nuclei, were 1.74 ± 0.67, 4.60 ± 0.32, and 2.87 ± 1.99, with the grand mean equal to 3.07 ± 1.44. This grand mean value was subtracted from nominal granule cell counts determined in Figure C-F, and the normalized adjusted values were plotted in Figure 2K.
